# Accurate genotyping of three major respiratory bacterial pathogens with ONT R10.4.1 long-read sequencing

**DOI:** 10.1101/2024.10.03.616467

**Authors:** Nora Zidane, Carla Rodrigues, Valérie Bouchez, Martin Rethoret-Pasty, Virginie Passet, Sylvain Brisse, Chiara Crestani

## Abstract

High-throughput massive parallel sequencing has significantly improved bacterial pathogen genomics, diagnostics, and epidemiology. Despite its high accuracy, short-read sequencing struggles with complete genome reconstruction and assembly of extrachromosomal elements such as plasmids. Long-read sequencing with Oxford Nanopore Technologies (ONT) presents an alternative that offers benefits like real-time sequencing and cost-efficiency, particularly useful in resource-limited settings. However, the higher error rates of ONT have so far limited its application in high-precision genomic typing. The recent release of ONT’s R10.4.1 chemistry, with significantly improved raw read accuracy (Q20+), offers a potential solution to this problem.

The aim of this study was to evaluate the performance of ONT’s latest chemistry for bacterial genomic typing against the gold standard Illumina technology, focusing on three respiratory pathogens of public health importance, *Klebsiella pneumoniae, Bordetella pertussis*, and *Corynebacterium diphtheriae*, and their related species. Using the Rapid Barcoding Kit V14, we generated and analyzed genome assemblies with different basecalling tools and models, at different simulated depths of coverage. ONT assemblies were compared to the Illumina reference for completeness and core genome multilocus sequence typing (cgMLST) accuracy (number of allelic mismatches).

Our results show that genomes obtained from raw data basecalled with Dorado (with both simplex and duplex reads) SUP v0.7.1, assembled with Flye, and with a minimum coverage depth of 30×, optimized the accuracy for all bacterial species tested. The error rates were consistently below 1% of each cgMLST scheme, indicating that ONT R10.4.1 data is suitable for high-resolution genomic typing applied to outbreak investigations and public health surveillance.

## Introduction

Whole genome sequencing has revolutionized the study of bacterial pathogens, emerging as crucial tool for molecular diagnostics and epidemiology, and as cornerstone of public health and clinical microbiology (Bagger et al., 2024; Doll et al., 2024; Revez et al., 2017). Over the past two decades, short-read sequencing technologies have dominated the research field and the market for molecular diagnostics and public health surveillance due to their high throughput and low error rates (Fox et al., 2014; Pfeiffer et al., 2018). This has provided scientists worldwide with high-resolution data for bacterial strain subtyping, which is indispensable for accurate and reliable public health surveillance. Today, bacterial isolate differentiation and outbreak investigation are mainly carried out using Single-Nucleotide-Polymorphism (SNP) analysis and gene-by-gene methods, including core genome multilocus sequence typing MLST (cgMLST) schemes. These schemes are available on curated databases like BIGSdb-Pasteur, PubMLST (Jolley et al., 2018; Jolley & Maiden, 2010), and EnteroBase (Zhou et al., 2020), facilitating standardized genomic typing, surveillance, and outbreak investigation of key pathogens (e.g., *Listeria monocytogenes*, and *Salmonella enterica*) from short-read data or draft assemblies.

However, reconstructing a complete bacterial genome *de novo* from short-read data is rarely possible due to complex, repetitive genomic regions such as insertion sequences (IS) and other repetitive elements (Ring et al., 2018). Short reads also struggle to reconstruct extra-chromosomal elements, such as plasmids (Arredondo-Alonso et al., 2017), making it difficult to map specific genes, like antimicrobial resistance (AMR) genes, to either the chromosome or mobile genetic elements. Additionally, second generation sequencing technologies like Illumina remain relatively expensive, both in terms of price *per* genome and acquisition cost of these sequencing platforms. These factors, along with limited portability, hinder their use in small labs and low- and middle-income settings.

Oxford Nanopore Technologies (ONT) sequencing overcomes several of these issues, including portability (e.g., MinION device), cost-efficiency (especially when multiplexing; Sanderson et al., 2024), and the ability to circularize chromosomes and plasmids due to its long-read nature (Lerminiaux et al., 2023). It also provides options for real-time sequencing and rapid library preparation, essential for quick outbreak responses (Wagner et al., 2023). However, ONT’s high error rate (Dohm et al., 2020; Jain et al., 2017) has so far limited its use in surveillance, as it impacts particularly gene-by-gene approaches like cgMLST. In fact, spurious SNPs introduced by sequencing errors can create artificial alleles, increasing allelic distances between isolates. As a result, most public bacterial genotyping databases do not currently accept ONT-only data.

Recent studies have attempted to benchmark ONT sequencing for bacterial genomic typing, showing variable but promising results depending on laboratory and bioinformatics factors (Lerminiaux et al., 2023; Sanderson et al., 2023, 2024; Soto-Serrano et al., 2024; Wagner et al., 2023), as well as on the pathogen (Linde et al., 2023; Sanderson et al., 2023). The new ONT chemistry R10.4.1 may address the high error rates of previous versions (e.g., R9), offering a declared raw read accuracy comparable to short-read technologies (>99%, Q20+). This study aimed to compare the performance of the existing gold standard short-read sequencing technology for bacterial genotyping (Illumina) with the latest ONT chemistry for cgMLST typing of three key respiratory pathogens, *Klebsiella pneumoniae, Corynebacterium diphtheriae* (the major agent of diphtheria), *Bordetella pertussis* (the agent of whooping cough) and their related species, using the Rapid Barcoding Kit V14.

## Materials and methods

### Bacterial isolates included in the study

Twenty-four isolates from 12 bacterial species were included in this study (Supplementary Dataset S1). Isolates belonged to the genera *Klebsiella* (in particular, to the *K. pneumoniae* Species Complex, or KpSC; genome sizes 4.7-6.3 Mbp), *Corynebacterium* (in particular, to the Corynebacteria of the *diphtheriae* Species Complex, or CdSC; genome sizes 2.2-2.9 Mbp), and *Bordetella (B. pertussis, B. parapertussis, B. holmesii* and *B. bronchiseptica;* genome sizes 3.3-5.6 Mbp). Isolates are either reference or type strains, or they were selected among the bacterial collections of our laboratory: i) the KpSC collection, ii) the French National Reference Centre (NRC) for CdSC collection, and iii) the French NRC for Whooping cough and other *Bordetella* infections collection.

### Isolate growth and DNA extraction

KpSC and CdSC were plated on Tryptic Soy Agar (TSA) and grown at 37 °C for 24h, and at 35–37 °C for 24-48 hours, respectively. *Bordetella* spp. isolates were grown at 36°C for 24 to 72 hours on Bordet-Gengou agar (Becton Dickinson, Le Pont de Claix, France) supplemented with 15% defibrinated horse blood (BioMérieux, Marcy l’Étoile, France) and subcultured in the same medium for 24 hours in standardized conditions, as previously described (Bouchez et al., 2018).

DNA extraction was performed on a Maxwell RSC Instrument (Promega, Madison, USA) with the Maxwell RSC Blood DNA Kit (Promega, Madison, Wisconsin, USA) following the manufacturer’s instructions.

### Library preparation and whole genome sequencing

Libraries for short-read sequencing were prepared and sequenced at the Mutualized Platform for Microbiology (P2M, Institut Pasteur) using the Nextera XT DNA library preparation kit (Illumina, San Diego, USA) on a NextSeq-500 or NextSeq 2000 apparatuses (Illumina, San Diego, USA) with a 2 × 150 nt paired-end protocol.

Libraries for long-read sequencing were prepared with the Rapid Barcoding Kit V14 (SQK-RBK114.24; Oxford Nanopore Technologies, Oxford, UK), and sequenced on three R10.4.1 flow cells (FLO-MIN114, one per pathogen) on a GridION machine for 72 hours. The minimal fragment length was set at 200 bp on the GridION software, v23.11.7. All libraries included a negative control barcode prepared with nuclease-free water.

### Long read basecalling and data processing

We tested different combinations of basecalling programs, basecalling models, data filtering and coverage to establish the best workflow possible (Figure 1). All the combinations used to generate the final assemblies can be found in Table 1, and all the bioinformatic commands can be found in the Supplementary information file.

**Table 1.**
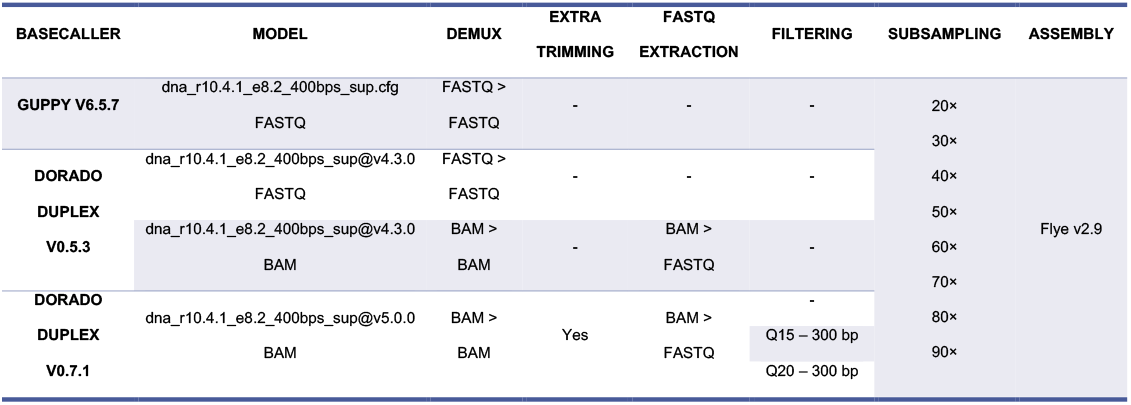
Different combinations of data processing used in this work, which generated long-read genome assemblies with different depths of coverage.

**Figure 1.**
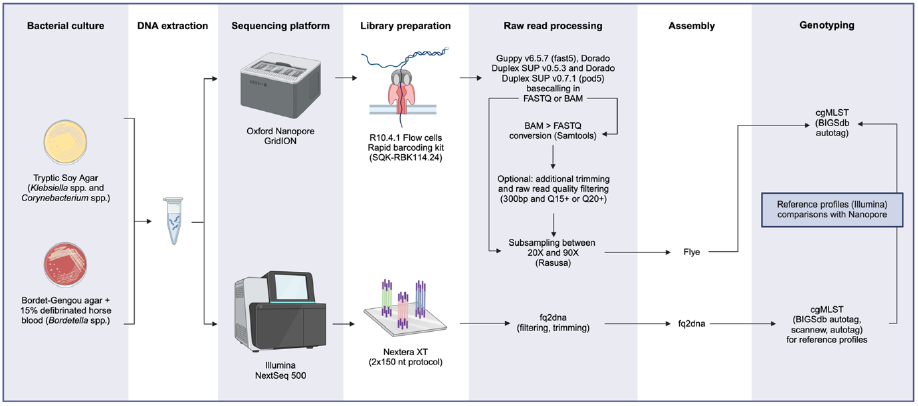
Graphical summary showing the experimental workflow followed in this study.

Three basecalling tools/versions were tested, but uniquely with super accurate (SUP) models, as these are known to lead to more accurate results (Lerminiaux et al., 2023). Data in fast5 format was basecalled and demultiplexed with Guppy v6.5.7 (https://community.nanoporetech.com). Data in pod5 format (converted from fast5 data with the pod5 v0.2.4 toolkit, https://github.com/nanoporetech/pod5-file-format) was processed with two versions of Dorado Duplex (basecalling both simplex and duplex reads at the same time; https://github.com/nanoporetech/dorado), v0.5.3 and v0.7.1 (Table 1). Data processed in a bam format was converted to fastq with Samtools v1.18 (Danecek et al., 2021).

Raw read quality was assessed with NanoStat v1.6.0 (De Coster et al., 2018), and data was plotted with python Seaborn v0.13.2 (Waskom, 2021).

To test the effect of raw read quality filtering on the final assembly, we generated different filtered read sets based on read length and read quality (minimum 300bp long and either minimum Q15 or Q20) with NanoFilt v2.8.0 (De Coster et al., 2018). This was done uniquely for data basecalled with Dorado Duplex v0.7.1, as not enough data with Q15+ scores was available for the other basecalled read sets (see the Results and discussion section). Long-reads were then subsampled with Rasusa v0.8.0 (Hall, 2022) to simulate different depths of coverage (from 20× to 90×, based on the maximum coverage possible per isolate).

### *De novo* assembly

Short-read sequence data was assembled with fq2dna v21.06 (https://gitlab.pasteur.fr/GIPhy/fq2dna). Subsampled long-read data sets were assembled with Flye v2.9 (Kolmogorov et al., 2019). We did not compare results from multiple assembly algorithms, as Flye has already been shown to lead to more complete and accurate genome assemblies compared to other tools (e.g., Unicycler, Raven; Lerminiaux et al., 2023). Variability in terms of average genome coverage was detected in the final subsampled assemblies (Figure S1), with most showing a coverage lower than the one used for subsampling, but higher than the lower ten (e.g., for 30× coverage used for subsampling, most assemblies had a coverage lower than 30× but higher than 20×). For this reason, results are given in a “greater than” format (e.g., for data subsampled at 30×, results are given as >20×). Assembly quality was checked with QUAST v5.0.2 (Gurevich et al., 2013).

### *Klebsiella* spp., *Corynebacterium* spp. and *Bordetella* spp. genomic typing

Genome assemblies generated with both short and long-read sequencing were uploaded to BIGSdb-Pasteur (https://bigsdb.pasteur.fr/) in their respective species databases. We defined cgMLST alleles on reference Illumina assemblies with the BIGSdb software (Jolley & Maiden, 2010), and subsequently tagged long-read assemblies for these alleles. The cgMLST schemes used for genotyping were: i) for KpSC isolates, the scgMLST629_S scheme (including 629 loci; Hennart et al., 2022); ii) for *C. diphtheriae* and *C. rouxii*, the *C. diphtheriae* cgMLST scheme (1,305 loci; Guglielmini et al., 2021), and for *C. ulcerans* the cgMLST_ulcerans scheme (1,628 loci; Crestani et al., 2024); iii) for *Bordetella* species (*B. pertussis, B. parapertussis, B. bronchiseptica*, and *B. holmesii)*, the cgMLST_genus scheme (1,415 loci; Bridel et al., 2022). Additionally, we used the cgMLST_pertussis scheme (2,038 loci; Bouchez et al., 2018) on *B. pertussis* genomes.

Minimum-spanning trees (MSTrees) based on cgMLST profiles of genome assemblies generated from Dorado Duplex SUP v0.7.1 data were constructed with GrapeTree (Zhou et al., 2018) for each genus.

In addition, Life Identification Numbers (LIN) codes using a gene-by-gene approach (Hennart et al., 2022; Palma et al., 2024) were assigned to KpSC assemblies.

### Allelic mismatch analysis and data visualization

The number of allelic mismatches between cgMLST profiles of Illumina vs ONT assemblies were computed with python pandas v1.4.3 (The pandas development team, 2020). Missing loci from short-read reference genomes were not considered for the analysis. The type of mismatches obtained by this comparison method include: i) spurious SNPs generating by chance alleles existing in the database; ii) spurious SNPs generating new artificial alleles (as no new alleles were defined on long-read assemblies, these would appear as missing data in the profile). The script used to compare cgMLST profiles can be found on GitHub (https://github.com/chcrestani/Comparison_cgMLST_profiles/).

All graphs were generated with python seaborn v0.13.2 (Waskom, 2021).

## Results and discussion

### Most raw reads generated with Dorado Duplex SUP v0.7.1 have Q20+ quality scores

Raw reads produced by basecalling with Guppy v6.5.7 had the lowest quality scores, with a median read quality of 15 across the three pathogens (Figure S2). Quality scores increased with Dorado and with its versions: from a median raw read quality between 19 and 20 with Dorado Duplex v0.5.3, to 21-23 with Dorado Duplex v0.7.1; for the latter, most of the data showed Q20+ quality scores (Figure S2). Scores of less than Q18 for Guppy and over Q20 for Dorado are consistent with previous work that also assessed performance of the Rapid Barcoding Kit V14 (Hall et al., 2024; Lerminiaux et al., 2023).

### Genome completeness

Illumina assemblies comprised between 14 and 295 contigs (Figure S3), with the *Bordetella* genus showing the highest average number of contigs (n=186). This is not surprising considering the high number of IS copies carried by *Bordetella* species, a known cause of genome fragmentation for *de novo* assembly of short-read data (Ring et al., 2018). *Klebsiella* genomes had an average of 53 contigs, and *Corynebacterium* genomes an average of 27.

ONT genomes consisted of one to eight contigs (Figure S3). None of the *Bordetella* isolates carried plasmids, and most ONT genomes were correctly assembled in a single circularized contig (96%, 295/307). The remaining *Bordetella* ONT assemblies (3.9%, 12/307) were fragmented in 2-6 contigs, likely a consequence of their low depths of coverage (10×-20×). For *Klebsiella* isolates, the expected number of contigs (corresponding to the chromosome and/or one or more plasmids) was detected in most ONT genome assemblies (6/8 isolates showing one main peak in Figure S4). For two isolates (SB11 and SB132) little consensus was observed among genomes in the assembly set (Figure S4), with most of the genome assemblies of SB11 showing loss of small plasmids. Among the *Corynebacterium* isolates, six had no plasmids and two (FRC1385 and FRC1356) carried one plasmid. Chromosomes and plasmids were correctly assembled in 77% of *Corynebacterium* ONT genomes (270/349; Figure S3), whereas they were too fragmented in 23% of cases (79/349, with a maximum of seven contigs).

Our data shows that the latest ONT chemistry, R10.4.1, with the Rapid Barcoding Kit V14, allows in most cases for complete reconstruction and circularization of both the chromosome and extra-chromosomal elements. This is particularly encouraging with regards to the genus *Bordetella*, for which assembly of short-read data has long presented limitations, as explained above. In addition, the reconstruction of extra-chromosomal elements is particularly useful in the case of bacteria carrying AMR or virulence plasmids, such as KpSC species. The observed loss of certain KpSC plasmids (especially of 9-12 kb) is not suspiring considering that some long-read assemblers, including Flye, are known to underperform in the reconstruction of small plasmids (Johnson et al., 2023). However, the overall plasmid loss was minimal across our assembly sets, and this can be mainly attributed to the choice of the Rapid barcoding kit over the Ligation kit, the latter being a known cause of underrepresentation of small plasmids in genome assemblies (Wick et al., 2021)

### Basecalling with Dorado Duplex SUP v0.7.1 allows for accurate genomic typing of the three pathogens

We analyzed the number of cgMLST mismatches after allele calling, compared to Illumina assembly calls used as reference. In most cases, the average number of mismatches per isolate did not differ significantly between data basecalled with Guppy v6.5.7 and with either Dorado Duplex version, as it was already low with Guppy-generated assemblies (Figure S5). However, in a few cases Guppy assemblies showed a non-negligible number of allelic mismatches, whereas Dorado Duplex v0.5.3 basecalling allowed to recover almost all the correct alleles (e.g., in *K. pneumoniae* isolate SB3928, in *C. diphtheriae* CIP100721^T^, and in *B. bronchiseptica* FR7093; Figure S5). No significant difference was observed between data basecalled directly in FASTQ format compared to data basecalled in BAM, which is Dorado’s default format. Two isolates (*K. variicola* subsp. *variicola* SB48 and *C. rouxii* FRC0190^T^) still showed a high number of mismatches in the assemblies generated with Dorado Duplex v0.5.3, even at high depths of coverage (between 116 and 243 for FRC0190^T^, and between 17 and 34 for SB48). Remarkably, Dorado Duplex v0.7.1 allowed to reduce this number to a level comparable to that of the other isolates belonging to the same genus (Figure S5).

Therefore, among the three basecalling tools and models used, Dorado Duplex v0.7.1 with the dna_r10.4.1_e8.2_400bps_sup@v5.0.0 model was much more accurate, likely as a result of the overall higher quality of the raw reads (Figure S2). Results reported from here onwards only refer to data generated with this model.

Allelic mismatches appear minimized at coverage depths >20× for KpSC and *Bordetella* spp. (Table 2, Figure 2), with an average number of mismatches >20× of 0.34/629 for the former (accounting for a 0.05% error rate of the scgMLST scheme) and 2.2/1,415 for the latter (0.16% error rate for the cgMLST_genus). A coverage depth of >30× was needed to minimize allelic mismatches for CdSC (Table 2), which resulted in an average number of allelic mismatches of 2.79. This corresponds to an error rate of 0.21% of the cgMLST diphtheriae scheme (1,305 loci), and of 0.17% of the cgMLST_ulcerans scheme (1,628 loci).

**Table 2.** Average number of mismatches and standard deviations (SD) at each depth of coverage detected in data basecalled with Dorado Duplex SUP v0.7.1. The means were calculated on all data regardless of quality filtering. For the isolates belonging to the Corynebacteria of the *diphtheriae* Species Complex, averages were calculated as the total number of mismatches regardless of the cgMLST scheme.

| Dorado Duplex v0.7.1 (unfiltered, Q15+ and Q20+) |  |  |  |
| --- | --- | --- | --- |
|  | *Klebsiella pneumoniae* Species Complex (scgMLST629_S) | *Corynebacteria* of the *diphtheriae* Species Complex (cgMLST diphtheriae or cgMLST_ulcerans) | *Bordetella* spp. (cgMLST_genus) |
| Coverage | Mean (SD) | Mean (SD) | Mean (SD) |
| >10× | 1.04 (±1.23) | 7.50 (±7.49) | 3.42 (±1.93) |
| >20× | 0.38 (±0.65) | 4.25 (±4.94) | 2.21 (±1.35) |
| >30× | 0.21 (±0.51) | 3.17 (±3.55) | 2.42 (±1.28) |
| >40× | 0.38 (±0.74) | 2.71 (±3.10) | 2.10 (±1.26) |
| >50× | 0.43 (±0.75) | 2.74 (±3.33) | 2.22 (±1.17) |
| >60× | 0.33 (±0.58) | 2.4 (±3.05) | 2.31 (±1.45) |
| >70× | 0.30 (±0.57) | 3.06 (±3.38) | 2.17 (±1.53) |
| >80× | 0.27 (±0.46) | 2.64 (±3.00) | 2.08 (±1.44) |

**Figure 2.**
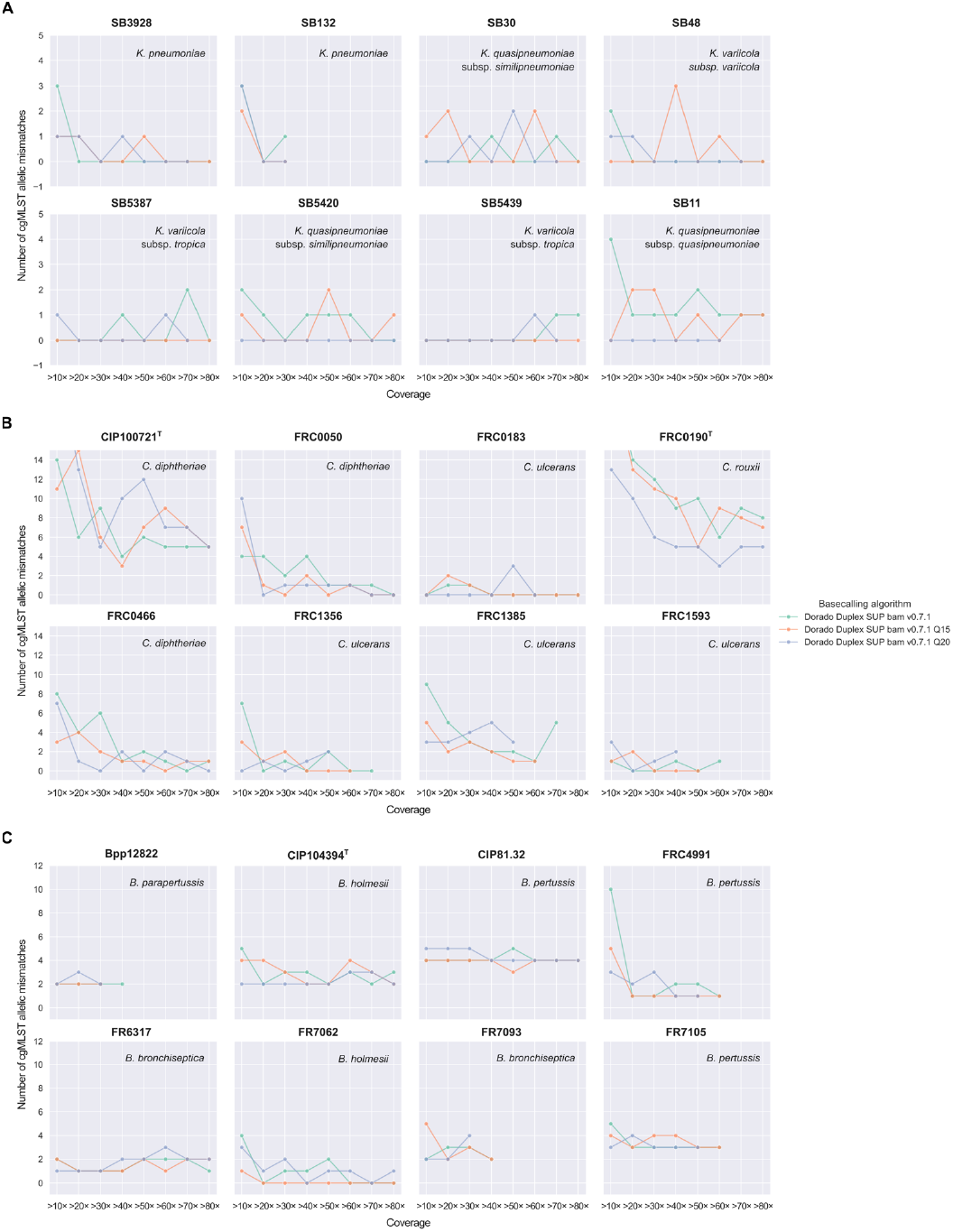
Total number of allelic mismatches between short-read assemblies generated with Illumina and long-read assemblies generated with Nanopore R10.4.1 sequencing from data basecalled with Dorado Duplex SUP v0.7.1. Mismatches are shown at different simulated depths of coverage for assemblies generated from unfiltered data, from data filtered with a minimum Phred quality score of 15 and of 20. A) Mismatches for isolates belonging to the *Klebsiella pneumoniae* Species Complex (KpSC); B) Mismatches for isolates belonging to the Corynebacteria of the *diphtheriae* Species Complex (CdSC); note that the y-axis was interrupted at fifteen allelic mismatches to improve readability of the graphs; C) Mismatches for *Bordetella* isolates, *including B. pertussis, B. parapertussis, B. holmesii* and *B. bronchiseptica*.

An error rate below 1% across all cgMLST schemes tested is reassuring, and it shows how it is nowadays possible to achieve high-resolution genomic typing of these three important respiratory pathogens with ONT R10.4.1 data. Additional steps of polishing (e.g., with Medaka) may be carried out when fast or high accuracy (HAC) models are used, as they have been shown to increase assembly quality (Arredondo-Alonso et al., 2017; Foster-Nyarko et al., 2023; Sanderson et al., 2023), but they do not appear necessary with SUP (Ryan Wick’s blog, 2023). We anticipate that new and improved versions of Dorado, together with the latest chemistry transition that was silently released by Nanopore during the first quarter of 2024 (new motor protein E8.2.1), should lead to yet less allelic mismatches. Based on ONT future developments, an even higher genome assembly quality could be attained thanks to an increased proportion of duplex data, which is known to have higher quality scores (>Q30 for SUP models) than simplex reads (Hall et al., 2024). In this work, duplex reads accounted for only 10-14% of the overall number of reads, which is consistent with previous findings (Soto-Serrano et al., 2024).

### Raw data quality filtering is not necessary but may be useful

We did not observe a significant effect of raw read quality filtering (Q15+ and Q20+) on the number of allelic mismatches in genomes from all three genera (Figure 2). However, the implementation of a filtering step with tools such as NanoFilt or Filtlong (https://github.com/rrwick/Filtlong) may be useful to reduce computational time or when computational resources are scarce. In addition, some authors have observed a decreased quality or completeness of genomes assembled with Flye at high depths of coverage (>50×-100×) (Ryan Wick’s blog, 2023, Thomas Cokelaer’s talk, 2023, Cokelaer et al., 2017). In our work, we observed either a stable or decreasing average number of mismatches with the increase of coverage depth (Figure S6). However, CdSC assemblies showed a progressively decreasing number of mismatches up to coverages >60×, with a slight increase >70×-80× (Figure S6). Therefore, although in our work data filtering based on quality did not have a significant effect on the total number of allelic mismatches, when too much data is generated, it may be useful to filter the raw reads based on quality to obtain a maximum average depth of coverage of 100×.

### ONT R10.4.1 sequencing can be used for rapid outbreak investigation and public health surveillance of KpSC, CdSC, and *B. pertussis*

Single-linkage classification and MSTrees with species-specific allelic mismatch thresholds may be used in public health for surveillance of multiple pathogens and for outbreak investigations. In addition, cgMLST-based LIN codes can be used to classify KpSC genomes and to detect outbreak strains (Palma et al., 2024). Here, we aimed to investigate the accuracy of ONT assemblies for these applications.

### *Klebsiella pneumoniae* Species Complex

When ignoring missing data, cgMLST profiles of ONT-assembled genomes had a maximum of four allelic mismatches compared to the Illumina reference (Figure 2, Figure S7). On the MSTree, the majority of KpSC profiles are part of the central genotype with the Illumina assembly (Figure S7). These profiles have identical alleles to the reference, and they are either complete, or they are missing one or more loci due to alleles that were not tagged because of spurious SNPs. In the latter case, they still cluster with the Illumina genotype in the MSTree because missing data is handled dynamically by GrapeTree, reducing the total number of loci considered in the pairwise distance calculation (i.e., GrapeTree computes the shortest possible connections between nodes to minimize the overall length of the tree). If these ONT assemblies had been scanned for new alleles, the currently incomplete profiles would appear as more distanced from the Illumina reference, which is why we do not recommend defining new alleles on ONT data at present.

LIN codes identical to that of the Illumina reference were detected for all cgMLST profiles of genomes with >20× coverage (Supplementary Dataset S1), with one exception: most of the ONT assemblies of SB30 had a LIN code that differed from the reference but that was identical among them. The difference was detected in bin number nine (second to last), which corresponds to a maximum of two allelic mismatches, and it was due to the presence in the ONT assemblies of an existing allele of a locus that was missing in the Illumina reference (allele 32; locus KP1_4024_S).

In previous work (Hennart et al., 2022), we observed that profiles of isolates involved in reported KpSC outbreaks generally differed by one or no allelic mismatches, with a maximum of five, and their LIN codes were either identical or differed in the last three bins. Here, our data shows how cgMLST profiles defined on ONT R10.4.1 genomes with >20× coverage can now be used for outbreak investigation of KpSC, as they perform similarly to Illumina-generated profiles with regards to allelic distances on MSTrees (GrapeTree) and to LIN codes.

### Corynebacteria of the *diphtheriae* Species Complex

In *C. diphtheriae* and *C. ulcerans*, all profiles were part of the central genotype with the Illumina reference on MSTrees generated with GrapeTree (Figure S8), despite the higher number of allelic mismatches detected compared to KpSC isolates (Figure 2, Table 2). This is explained by the fact that, whereas in KpSC some spurious SNPs in ONT assemblies generated random existing alleles (isolates appearing as distanced of 1-4 mismatches compared to the Illumina genotype; Figure S7), in CdSC the spurious SNPs did not lead to any existing allele being called.

A threshold of 25 allelic mismatches for single linkage groups was identified in previous work (Crestani et al., 2024; Guglielmini et al., 2021) as the maximum observed for known clusters of infection, and hence to define genetic clusters in both *C. diphtheriae* and *C. ulcerans*. Based on our analyses, ONT R10.4.1-generated cgMLST profiles defined by tagging existing alleles performs equally to Illumina data when classifying isolates into genetic clusters.

### *Bordetella pertussis* and other *Bordetella* species

For the genus *Bordetella* (using the cgMLST_genus scheme), the central GrapeTree genotype included the Illumina reference and most ONT genomes for *B. holmesii* (Figure 3) and *B. bronchiseptica* II. For the remaining isolates, including *B. pertussis, B. parapertussis and B. bronchiseptica* I-4, most ONT assemblies constitute the central genotype, and they all show one to three identical allelic mismatches compared to the gold standard Illumina (Figure 3). Most of these systematic mismatches (9/12) were artifacts: they were a result of two alleles of the same locus being present in the ONT assemblies (e.g., alleles “10;21” of locus BORD004759 in FR7093) and only one allele being called in the Illumina genome (e.g., allele 21 of locus BORD004759 in FR7093). For these double loci, one of the two alleles always corresponds to the Illumina one (Table S1). The detection of duplicated genes is a direct consequence of ONT genomes being more complete than Illumina assemblies, as described above. Therefore, when investigating outbreaks of *Bordetella* with the cgMLST_genus scheme, it is important to keep in mind that allelic distances between isolates could be overestimated.

**Figure 3.**
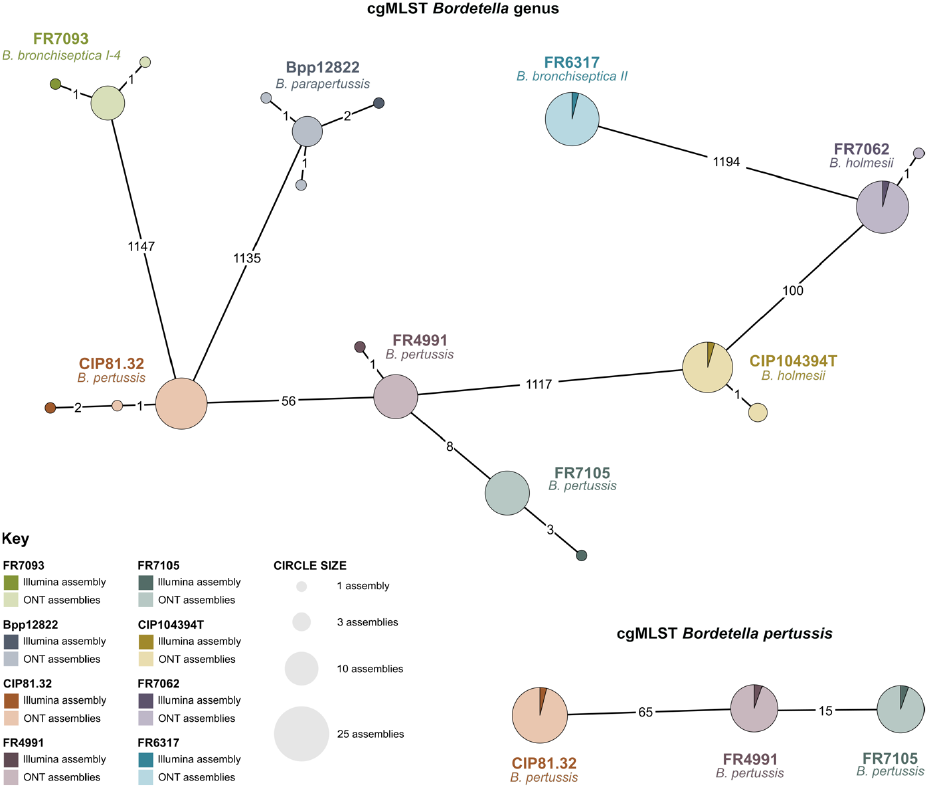
Minimum Spanning Trees of *Bordetella pertussis* and other *Bordetella* species investigated in this work (computed with GrapeTree). Core genome multilocus sequence typing (cgMLST) profiles used for pairwise comparisons in these trees were generated from i) Illumina genome assemblies (dark triangles); ii) Oxford Nanopore Technology (ONT) genome assembly, whose raw data was basecalled with Dorado Duplex SUP v0.7.1 (lighter colors). The upper tree was generated from cgMLST profiles of the cgMLST_genus scheme, whereas the lower tree from cgMLST profiles of the cgMLST_pertussis scheme (uniquely applied to *B. pertussis* genomes).

With regards to *B. pertussis*, we also investigated the performance of ONT R10.4.1 on the cgMLST scheme dedicated to this species. In this case, an average of 0.38 allelic mismatches was found at coverages >20×, which represents a very low error rate compared to the total number of loci of this scheme (n=2,038, 0.02%). Similarly to corynebacteria, these mismatches all corresponded to missing data, as can be observed in the MSTree (Figure 3), where all profiles from ONT assemblies are grouped in the central genotype with the Illumina reference.

ONT assemblies generated with the (best) parameters identified in this work appear therefore reliable to define recent chains of transmission with the *B. pertussis* scheme (current threshold of 3-4 allelic mismatches). Based on the issues caused by gene duplications with the cgMLST_genus scheme, we recommend using the cgMLST_pertussis one when investigating epidemics of *B. pertussis*.

## Conclusions

Public bacterial genome databases for bacterial strain taxonomy have so far not accepted genomes generated with previously existing ONT sequencing chemistries (e.g., R9.4.1) due to higher error rates compared to Illumina. This study evaluates the use of ONT R10.4.1 chemistry with the Rapid Barcoding Kit V14 for fast, high-resolution genomic typing of three respiratory pathogens (*K. pneumoniae, C. diphtheriae*, and *B. pertussis*) and related species curated on the BIGSdb-Pasteur database. The Rapid Barcoding kit was chosen for its cost-effectiveness, simplicity, and minimal laboratory requirements, making it ideal for low-resource and emergency settings (e.g., mobile diagnostic and sequencing laboratories). Despite earlier versions having higher error rates compared to the Native Barcoding Kit, recent studies suggest that assemblies generated with the Rapid Barcoding kit V14 are comparable to Illumina data (Sanderson et al., 2024).

Our data show that ONT genome assemblies of the three pathogens tested can be used for genomic strain typing if generated with the following workflow: library preparation and sequencing with the Rapid Barcoding Kit V14 on R10.4.1 flow cells, Dorado Duplex basecalling (v0.7.1 or above, and the latest SUP model available), Flye assembly (v2.9 or higher), and a minimum coverage of 30×. Other basecalling models, such as fast and HAC (not tested in this study), have been shown to lead to less accurate results (Lerminiaux et al., 2023). Thus, if computational resources are available, SUP models are to be preferred. Alternatively, polishing genomes generated with fast or HAC models with tools such as Medaka could be considered, as that has been shown to improve the final genome assembly quality (Arredondo-Alonso et al., 2017; Foster-Nyarko et al., 2023; Sanderson et al., 2023).

Though we currently advise to use ONT-generated genomes only for tagging existing cgMLST alleles and not for defining new ones due possible to spurious SNPs, this method still offers sufficient resolution for outbreak investigations and classifications (e.g., single-linkage clustering, MSTrees, and LIN code classification). The ability to perform precise genotyping with a low-cost, portable sequencing technology, such as ONT, represents a significant advance. It is also highly timely, considering the current epidemiological situation with i) rising cases of whooping cough in multiple world regions in 2024 (Fu et al., 2023; Pan American Health Organization, 2024; Rodrigues et al., 2024; ECDC, 2024), ii) one of the largest diphtheria outbreaks of recent times in West Africa (Balakrishnan, 2024; Samarasekera, 2024; WHO, 2024) and resurgence in Europe (Hoefer et al., 2023) and iii) the rising importance of multidrug-resistant infections caused by *K. pneumoniae* (Antimicrobial Resistance Collaborators, 2022; WHO, 2024). With ongoing technological advancements, and pending efficient procurement solutions, ONT could soon play a crucial role in global pathogen surveillance and outbreak response, including in low-resource settings.

## Data access

Genome assemblies (Illumina and ONT) can be downloaded from three projects in bigsdb.pasteur.fr (https://bigsdb.pasteur.fr/klebsiella/, project ID: 142; https://bigsdb.pasteur.fr/diphtheria/, project ID: 36; https://bigsdb.pasteur.fr/bordetella/, project ID: 50). All raw data generated in this study have been submitted to the NCBI BIoProject database, BioProject ID PRJNA1166325 for Illumina data, and Bioproject ID PRJNA1167647 for ONT data.

## Acknowledgements

We thank the Biomics Platform at Institut Pasteur for sharing their GridION machine, and in particular Chloé Baum for her support. We also thank the Mutualized Platform for Microbiology (P2M) for sequencing isolates using Illumina technology. This work used the computational and storage services provided by the IT Department at Institut Pasteur.

## Authors contributions

Conceptualization, Methodology and Visualization: CC; Data Curation, Validation, Formal Analysis: CC, CR, VB, MRP; Experimental work: NZ, VP; Writing original draft: CC; Writing review & editing: CC, CR, VB, SB; Resources and Funding Acquisition: SB.

