## Supplementary information for "Accurate genotyping of three major respiratory bacterial pathogens with ONT R10.4.1 long-read sequencing"

**Contents**

Supplementary figures S1-S8……………………………………………………………….p. 2-9
Supplementary table S1……………………………………………………………………p. 10
Bioinforamtic commands…………………………………………………………………...p. 11

### Supplementary figures

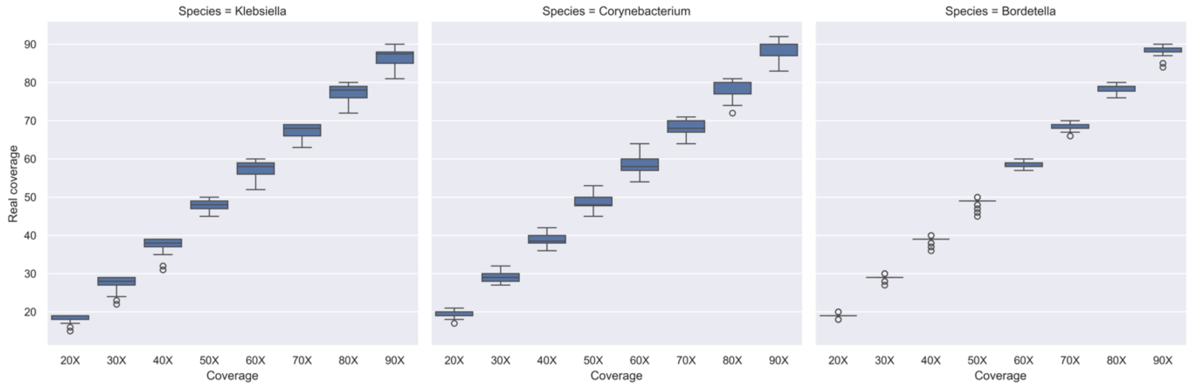


Figure S1. Box plots showing the average genome coverage (x-axis) used for subsampling with Rasusa and the real average genome coverage obtained in the final assemblies (y-axis). Most assemblies show a coverage lower than the one used for subsampling, but higher than the lower ten (e.g., for 30× coverage used for subsampling, most assemblies had a coverage lower than 30× but higher than 20×).

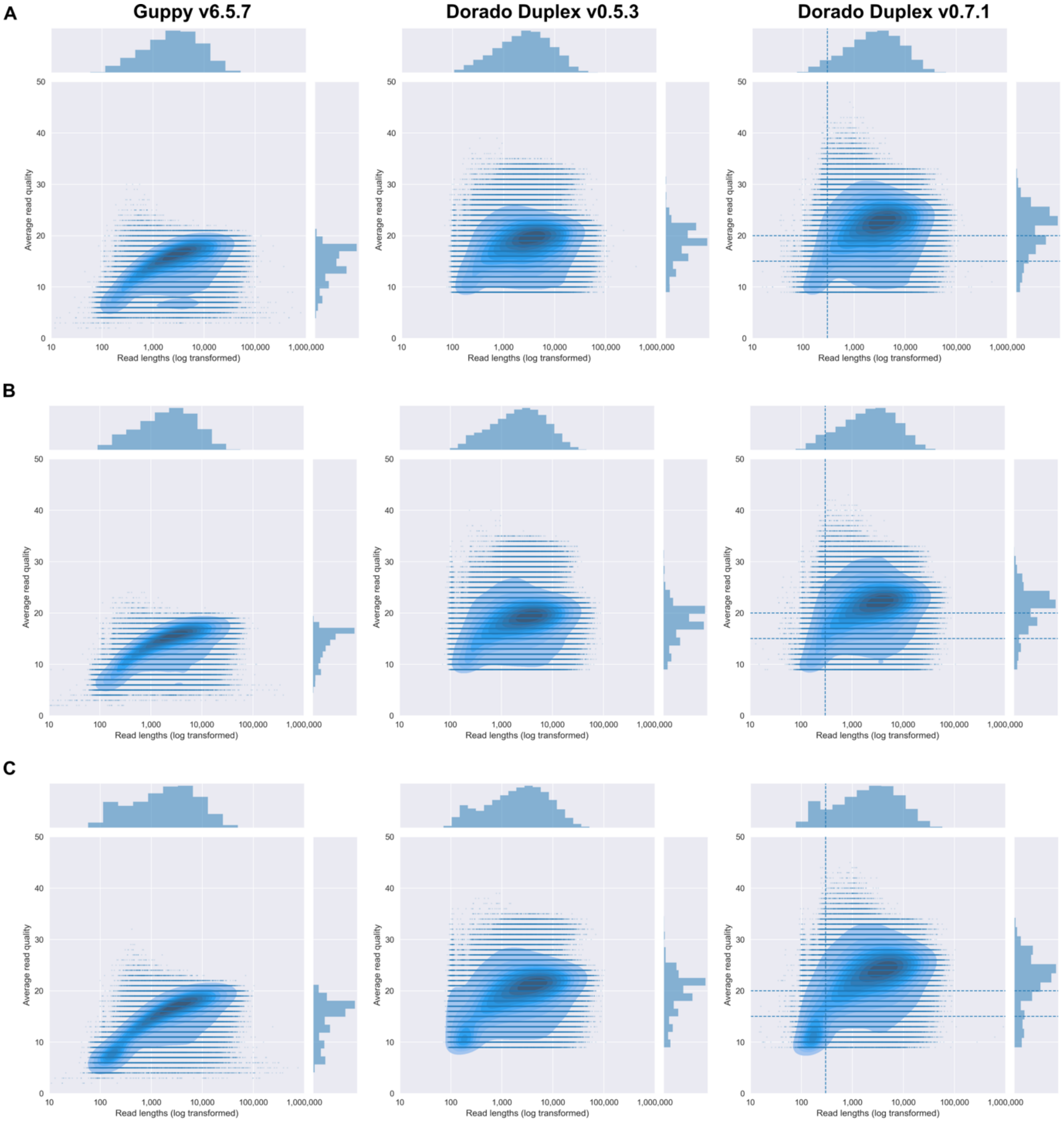


Figure S2. Raw read data quality and lengths with three basecalling algorithms, Guppy v6.5.7 SUP, Dorado Duplex v0.5.3 SUP and Dorado Duplex v0.7.1 SUP. Read length has been log transformed to allow better visualisation, but the x-axis indicates the actual read length. A) Results for the *Klebsiella pneumoniae* Species Complex library; B) Results for the Corynebacteria of the *diphtheriae* Species Complex library; C) Results for the *Bordetella* library.

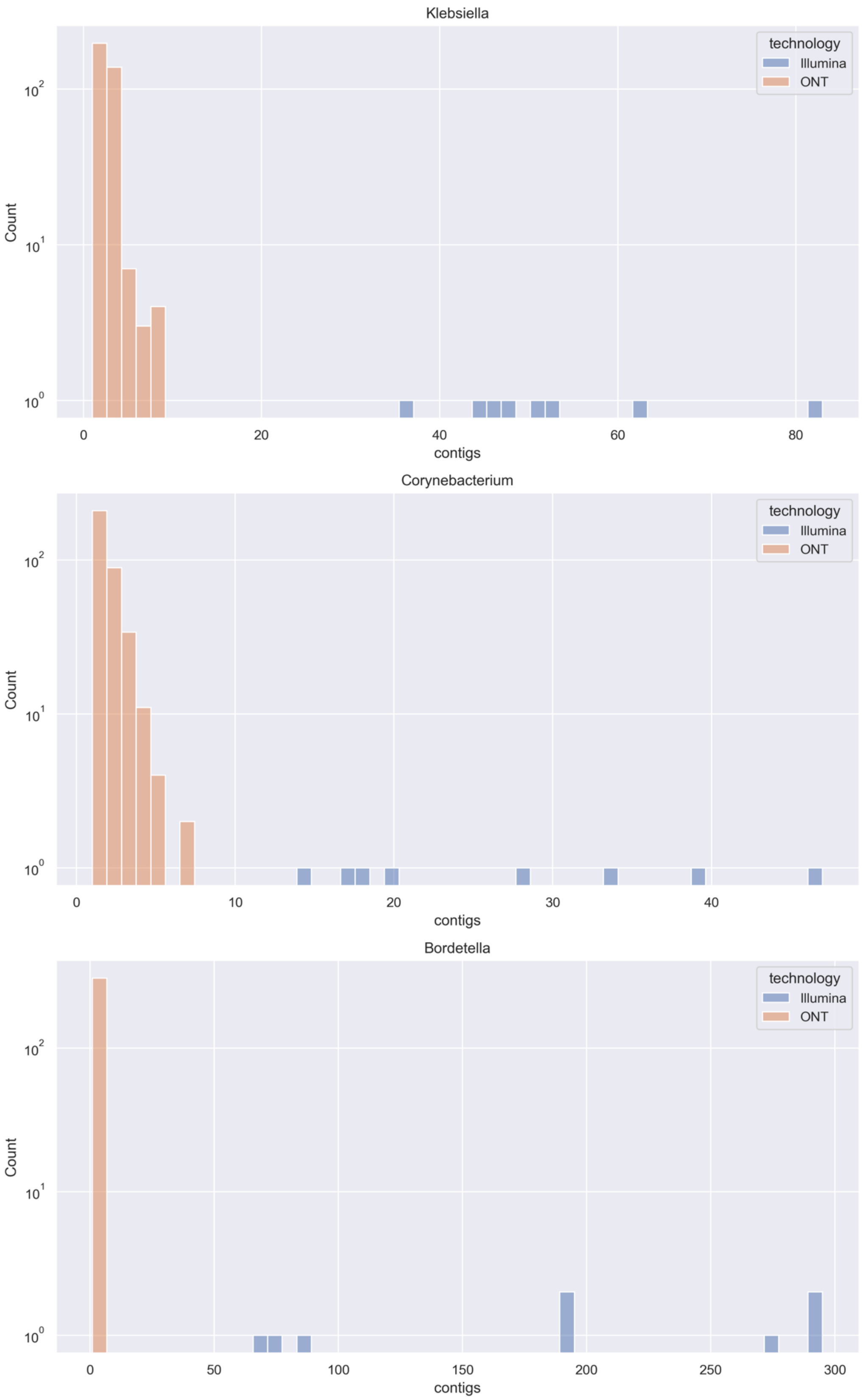


Figure S3. Distribution plots of the number of contigs detected in genomes of *Klebsiella, Corynebacterium* and *Bordetella* species, that were either generated with Illumina or Oxford Nanopore Technology (ONT). The y-axis was log-transformed.

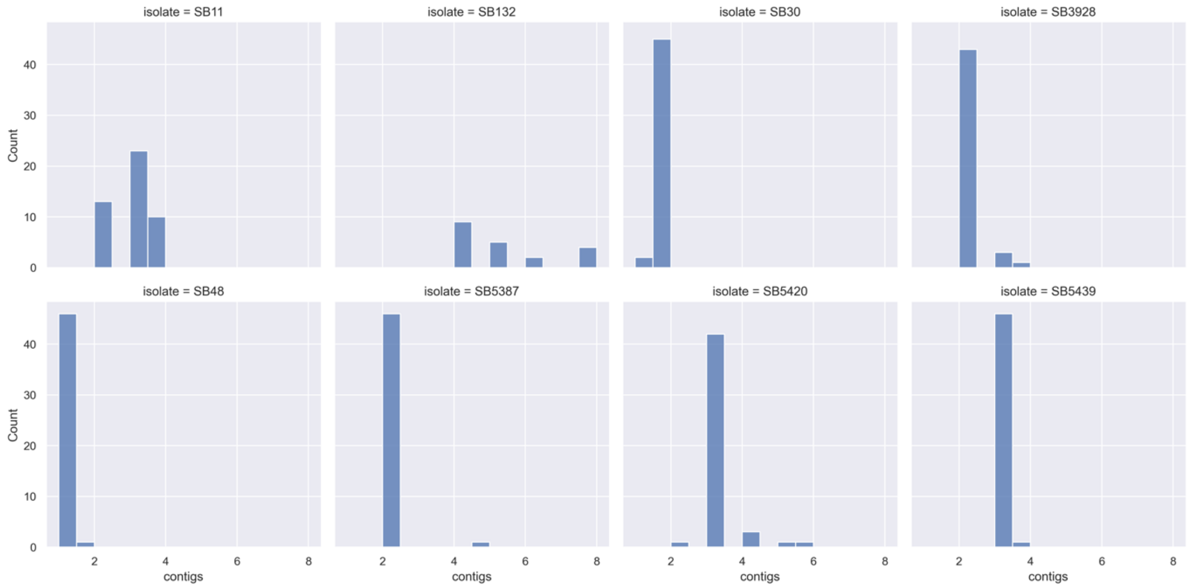


Figure S4. Number of contigs detected in *Klebsiella pneumoniae* Species Complex assemblies obtained with Oxford Nanopore Technology (ONT). In most cases, the expected number of contigs, corresponding to the chromosome and/or one or more plasmids, was detected (6/8 isolates showing one main peak). For two isolates (SB11 and SB132) a lot of uncertainty can be observed: in the case of SB11, which carries three plasmids (151kbp, 12kbp, 9kbp), some genome assemblies correctly identified all of them (four fragments, including the chromosome), but most of the genomes lost one or both smaller plasmids (2-3 fragments). For SB132, most of the genome assemblies have a closed chromosome and two circularized plasmids (105kbp and 96kbp), with a few genomes showing one (44kbp) or more additional linear contigs, which map to *Klebsiella* plasmids in NCBI. In a minority of assemblies of SB30 and SB5420, plasmids (77kbp and 48kbp) were lost.

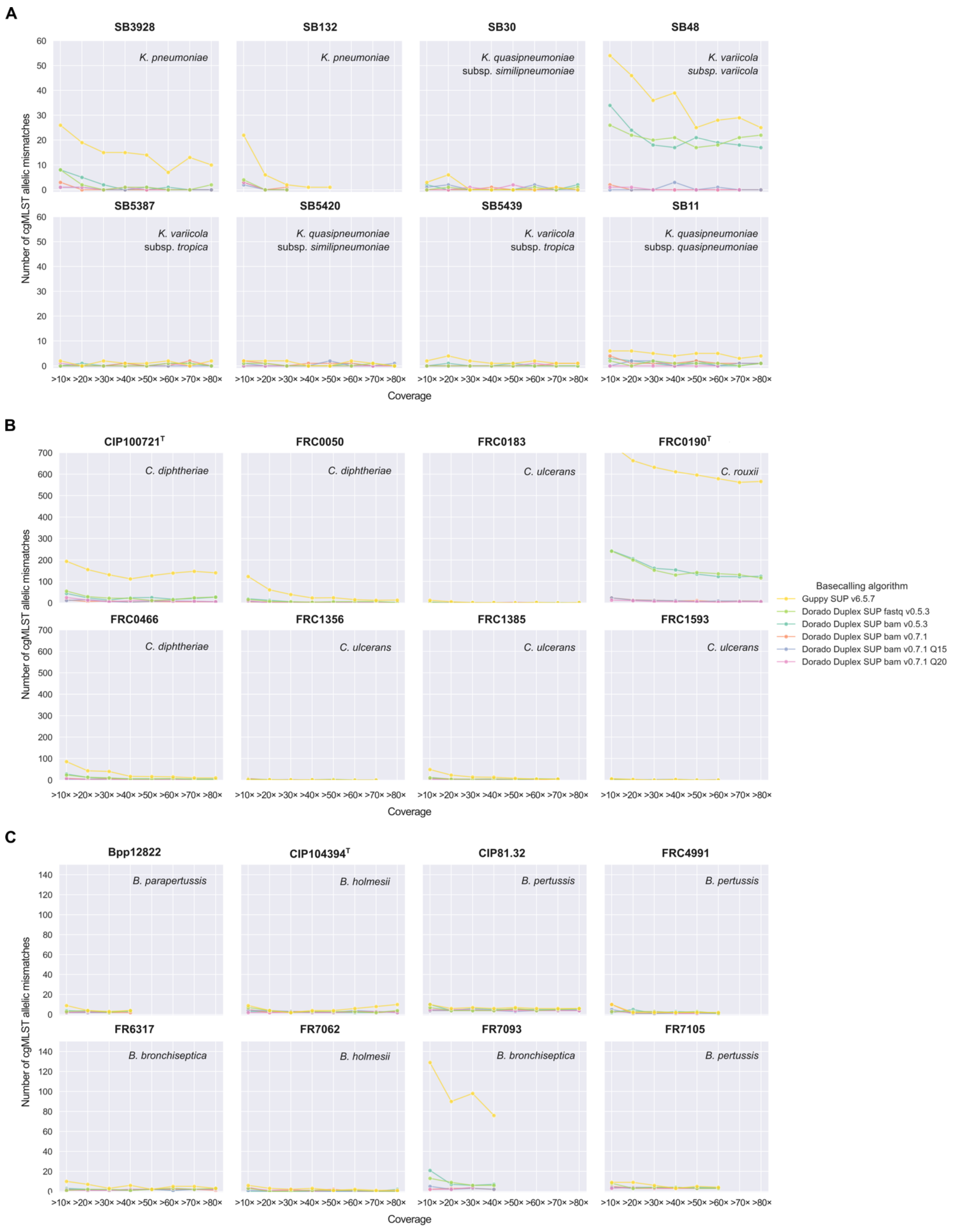


Figure S5. Total number of mismatches between short-read assemblies generated with Illumina and long-read assemblies generated with Nanopore R10.4.1 sequencing, at different depths of coverage. A) Mismatches for isolates belonging to the *Klebsiella pneumoniae* Species Complex (KpSC); B) Mismatches for isolates belonging to Corynebacteria of the *diphtheriae* Species Complex (CdSC); C) Mismatches for *Bordetella* isolates, including *B. pertussis, B. parapertussis, B. holmesii* and *B. bronchiseptica.*

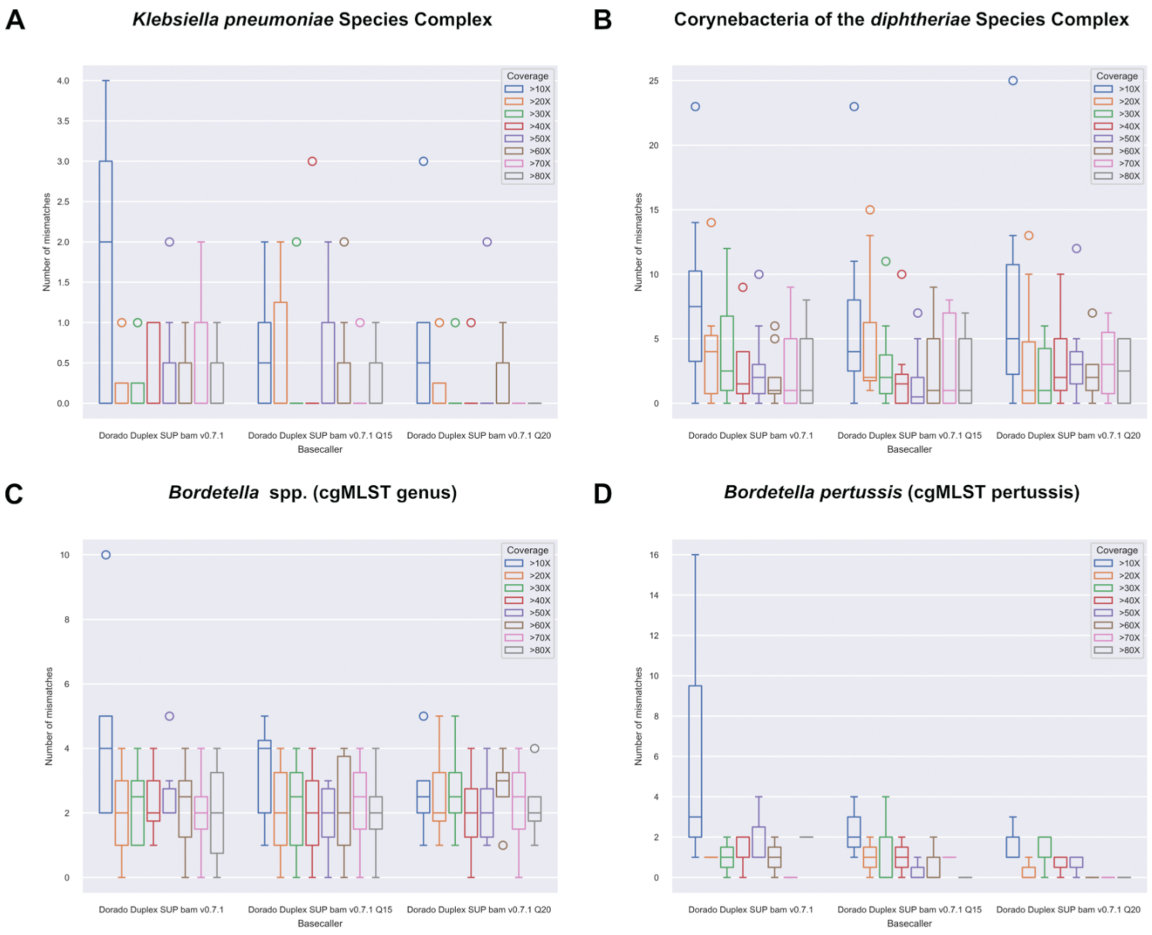


Figure S6. Boxplots for the number of mismatches per genome of data generated with Dorado Duplex SUP v0.7.1. A) *Klebsiella pneumoniae* Species Complex (scgMLST629_S scheme); B) Corynebacteria of the *diphtheriae* Species Complex (cgMLST and cgMLST_ulcerans scheme); C) *Bordetella* isolates (data from the cgMLST_genus scheme); D) *Bordetella pertussis* (data from the cgMLST_pertussis).

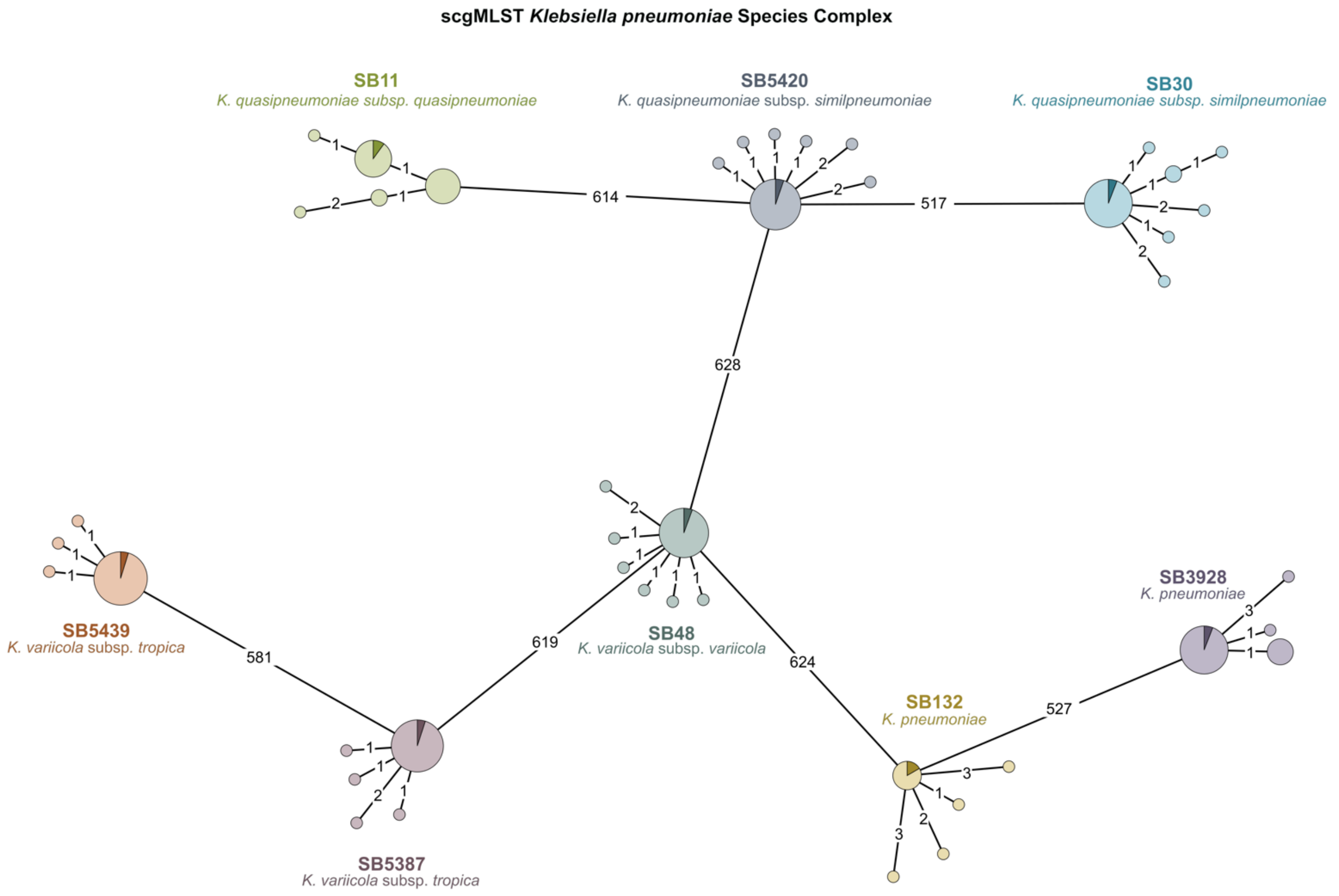


Figure S7. Minimum Spanning Tree of *Klebsiella pneumoniae* Species Complex (KpSC). Core genome multilocus sequence typing (cgMLST) profiles used for pairwise comparisons in these trees were generated from i) Illumina data (dark triangles); ii) Oxford Nanopore Technology (ONT) data from the basecaller Dorado Duplex SUP v0.7.1 (lighter colors). The tree was generated from cgMLST profiles of the scgMLST629_S scheme.

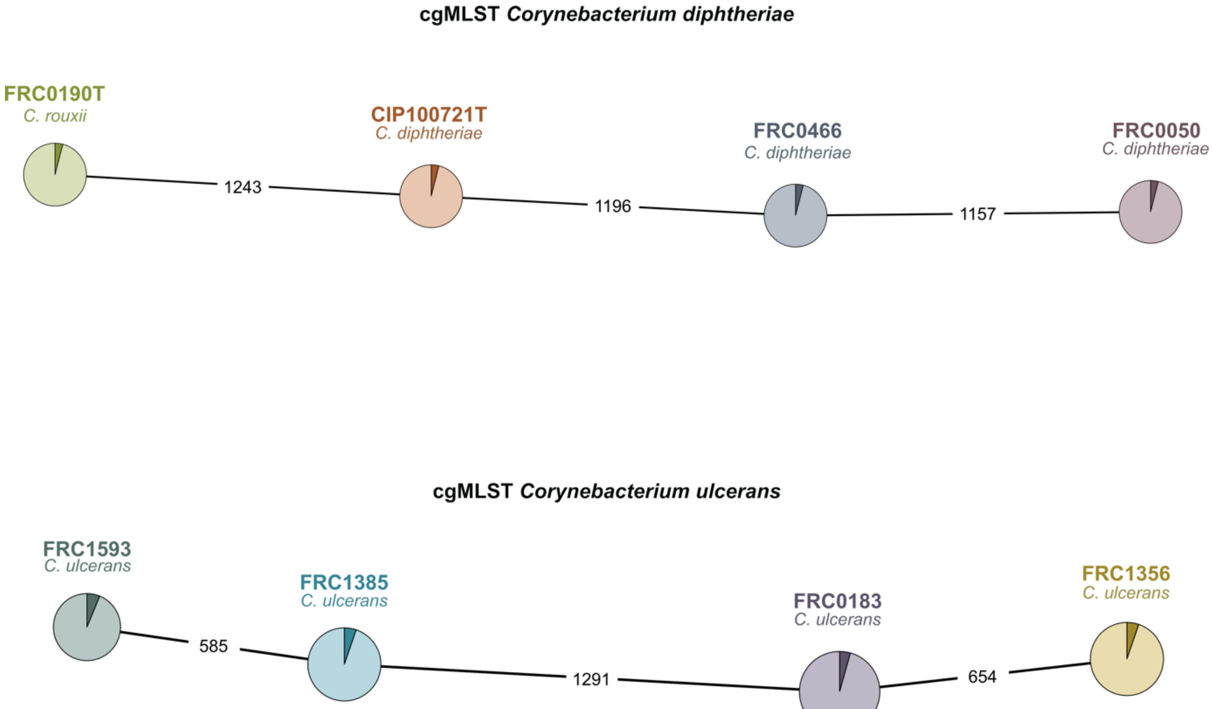


Figure S8. Minimum Spanning Tree of Corynebacteria of the *diphtheriae* Species Complex (CdSC). Core genome multilocus sequence typing (cgMLST) profiles of genome assemblies displayed in the tree were generated from i) Illumina data (dark triangles); ii) Oxford Nanopore Technology (ONT) data from the basecaller Dorado Duplex SUP v0.7.1 (lighter colors). The upper tree was generated from cgMLST profiles of the cgMLST diphtheriae scheme (*C. diphtheriae* and *C. rouxii*), whereas the lower tree from cgMLST profiles of the cgMLST_ulcerans scheme (uniquely applied to *C. ulcerans* genomes).

### Supplementary table

Table S1. Gene duplications in *Bordetella* species can lead to more than one allele of the same locus being detected and called (result shown as “allele1;allele2”). When comparing these allele calls with genomes in which only one allele was detected (mostly due to genome incompleteness) BIGSdb and GrapeTree will consider them as a mismatch, even if the one allele call matches one of the other two (e.g. allele 21 vs 10;21 of locus BORD004759 will be considered a mismatch). In most cases, Oxford Nanopore Technology (ONT) assemblies were more complete, and led to the detection of multiple alleles of the same locus. The only exception was for isolate FR7093, in which the Illumina assembly showed two alleles of the same locus (alleles 10;23, locus BORD000926), with most of the ONT assemblies giving the same result except for one genome assembly (FR7093_0.7.1_Q15_40X) that carried only one of the two alleles.

<table>
<tr><th>Species</th><th>Isolate</th><th>Locus</th><th>Illumina allele number</th><th>ONT allele number<br>(n of assemblies)</th></tr>
<tr><td rowspan="2">B. bronchiseptica I-4</td><td rowspan="2">FR7093</td><td>BORD004759</td><td>21</td><td>10;21 (11/11)</td></tr>
<tr><td>BORD000926</td><td>10;23</td><td>23 (1/11)</td></tr>
<tr><td rowspan="2">B. parapertussis</td><td rowspan="2">Bpp12822</td><td>BORD001951</td><td>21</td><td>3 (10/10)</td></tr>
<tr><td>BORD000926</td><td>33</td><td>3;33 (8/10)</td></tr>
<tr><td rowspan="8">B. pertussis</td><td rowspan="4">CIP81.32</td><td>BORD002924</td><td>6</td><td>5;6 (24/24)</td></tr>
<tr><td>BORD000926</td><td>5</td><td>4;5 (23/24)</td></tr>
<tr><td>BORD000877</td><td>4</td><td>4;21 (24/24)</td></tr>
<tr><td>BORD004953</td><td>7</td><td>3;7 (24/24)</td></tr>
<tr><td>FR4991</td><td>BORD000926</td><td>5</td><td>4;5 (17/17)</td></tr>
<tr><td rowspan="3">FR7105</td><td>BORD003456</td><td>37</td><td>4 (17/17)</td></tr>
<tr><td>BORD004760</td><td>36</td><td>4 (17/17)</td></tr>
<tr><td>BORD000926</td><td>5</td><td>4;5 (17/17)</td></tr>
</table>

### Bioinformatic commands

**Basecalling with Guppy**
guppy_basecaller --input_path ./fast5 --save_path guppy_basecaller_sup --min_qscore 7 --recursive -x 'cuda:0' --num_callers 4 --gpu_runners_per_device 8 --disable_pings -c dna_r10.4.1_e8.2_400bps_sup.cfg --compress_fastq

**Demultiplexing with Guppy**
guppy_barcoder -i ./pass -s guppy_barcoding_sup --trim_adapters

**Fast5 to pod5 conversion**
pod5 convert fast5 *.fast5 --output converted.pod5

**Basecalling with Dorado Duplex**

1. **On FASTQ files**
   dorado duplex ${DORADO_MODELS}/dna_r10.4.1_e8.2_400bps_sup@v5.0.0 --min-qscore 9 --emit-fastq converted.pod5 > dorado_sup_duplex_fastq.fastq
2. **On BAM files**
   dorado duplex ${DORADO_MODELS}/dna_r10.4.1_e8.2_400bps_sup@v5.0.0 --min-qscore 9 converted.pod5 > dorado_sup_duplex_bam.bam

**Demultiplexing with Dorado**

1. **On FASTQ files**
   dorado demux --kit-name SQK-RBK114-24 --output-dir dorado_demux_sup_duplex_fastq --emit-fastq dorado_sup_duplex_fastq.fastq
2. **On BAM files**
   dorado demux --kit-name SQK-RBK114-24 --output-dir dorado_demux_sup_duplex_bam dorado_sup_duplex_bam.bam

**Converting Dorado generated BAM files into FASTQ**
samtools bam2fq *.bam > *.fastq

**Obtain sequencing summary statistic from Dorado basecalled data**
dorado summary *.bam > summary_bam.tsv

**Read quality filtering (optional step)**
gunzip -c barcode*/*.fastq.gz | NanoFilt -q 15 -l 300 | gzip > barcode*/Q15_fil_reads.fastq.gz
gunzip -c barcode*/*.fastq.gz | NanoFilt -q 20 -l 300 | gzip > barcode*/Q20_fil_reads.fastq.gz

**Rasusa subsampling**
rasusa --input SQK-RBK114-24_$isolate.bam.fastq --coverage 20 --genome-size $size mb -o $isolate_20X.fastq.gz
rasusa --input SQK-RBK114-24_$isolate.bam.fastq --coverage 30 --genome-size $size mb -o $isolate_30X.fastq.gz
rasusa --input SQK-RBK114-24_$isolate.bam.fastq --coverage 40 --genome-size $size mb -o $isolate_40X.fastq.gz
rasusa --input SQK-RBK114-24_$isolate.bam.fastq --coverage 50 --genome-size $size mb -o $isolate_50X.fastq.g
rasusa --input SQK-RBK114-24_$isolate.bam.fastq --coverage 60 --genome-size $size mb -o $isolate_60X.fastq.gz
rasusa --input SQK-RBK114-24_$isolate.bam.fastq --coverage 70 --genome-size $size mb -o $isolate_70X.fastq.gz
rasusa --input SQK-RBK114-24_$isolate.bam.fastq --coverage 80 --genome-size $size mb -o $isolate_80X.fastq.gz
rasusa --input SQK-RBK114-24_$isolate.bam.fastq --coverage 90 --genome-size $size mb -o $isolate_90X.fastq.gz

**Flye assembly**
flye --nano-raw *.fastq.gz --genome-size $size m --out-dir $isolate_flye/ --threads 32
